# *Bmal1*-knockout mice exhibit reduced cocaine-seeking behaviour and cognitive impairments

**DOI:** 10.1101/2022.04.01.486740

**Authors:** Adriana Castro-Zavala, Laia Alegre-Zurano, Lídia Cantacorps, Ines Gallego-Landin, Patrick-S. Welz, Salvador A. Benitah, Olga Valverde

## Abstract

Brain and Muscle Arnt-like Protein 1 (BMAL1) is an essential component of the molecular clock underlying circadian rhythmicity. Recently, its function has also been associated with alterations in mood, and reward processing. We investigated the behavioural and neurobiological impact of *Bmal1* gene deletion in mice, as well as how these alterations affect rewarding effects of cocaine. Additionally, key clock genes and components of the dopamine system were assessed in several brain areas. Our results evidence behavioural alterations in *Bmal1-KO* mice including changes in locomotor activity with impaired habituation to environments as well as short term memory and social recognition impairments. In addition, *Bmal1-KO* mice experienced reduced cocaine-induced sensitization and rewarding effects of cocaine as well as reduced cocaine-seeking behaviour. Furthermore, *Bmal1* deletion influenced the expression of other clock-related genes in the mPFC and striatum as well as alterations in the expression of dopaminergic elements. Overall, the present article offers a novel and extensive characterization of *Bmal1-KO* animals. We suggest that reduced cocaine’s rewarding effects in these mutant mice might be related to *Bmal1* role as an expression regulator of MAO and TH, two essential enzymes involved in dopamine metabolism.

## INTRODUCTION

Circadian rhythms are endogenous oscillations lasting around 24 hours that synchronise physiology and metabolism with the external light-dark cycle. The suprachiasmatic nucleus (SCN) along with the numerous peripheral clocks distributed throughout the different tissues represents the core circadian machinery^1,2^. In this sense, the so-called clock genes maintain feedback loops that regulate the rhythmicity of biological processes.

In mammalian cells, brain and muscle ARNT-like protein 1 (*Bmal1*) heterodimerizes with circadian locomotor output cycles kaput (CLOCK) and drives the transcription of the gene families period (*Per*) and cryptochrome (*Cry*). PER-CRY protein complexes translocate into the nucleus and accumulate over time to repress their expression via suppression of CLOCK-BMAL1 activity. The subsequent degradation of PER and CRY proteins leads to reinitiating the cycle^3–5^. Additionally, proteins including nuclear receptors RORα and REV-ERB also activate and suppress *Bmal1* transcription, respectively, conferring additional stabilization to this molecular system ^6,7^.

Unlike most clock genes^8^, *Bmal1* is a non-redundant component of the circadian pacemaker and *Bmal1* knockout (KO) is the only single-gene deletion that profoundly impairs circadian rhythmicity^9^. Constitutive *Bmal1-KO* mice display reduced lifespan and early ageing phenotype, which is related to increased levels of reactive oxygen species^10^. Moreover, these mice also present reduced body weight and abnormal locomotor activity^11,12^.

Disruption of circadian rhythmicity is associated with the development of psychiatric disorders^13^. Data suggest a strong bidirectional relationship between circadian machinery and motivation processing^14–17^. While sleep disturbances are reported by drug users^18^, impairments in circadian rhythms impact motivational function and could lead to drug addiction^15,17,19^. The fact that dopaminergic activity displays circadian rhythmicity evidences the tight interplay between these systems^20,21^. Particularly, dopamine synthesis and degradation are under circadian control^22^. REV-ERBα represses tyrosine hydroxylase (TH) promoter^23^, while *Bmal1* directly regulates murine monoamine oxidase A (MAO-A) transcription^24^. Moreover, dopamine receptors 2 (D2R) and 3 (D3R) exhibit diurnal rhythm-dependent variations^25^, the latter being regulated by RORα and REV-ERB in the ventral striatum (STR)^26^.

Preclinical studies have evidenced that cocaine exposure alters clock genes^27–29^. However, whether disruption of circadian rhythms can influence subsequent cocaine-related responses has not yet been fully addressed. Mice lacking a functional *Clock* gene display increased cocaine reward as per the conditioned place preference (CPP) paradigm^30^. Similarly, deletion of the circadian transcription factor neuronal PAS domain protein 2 (NPAS2) increases cocaine self-administration (SA)^31^. However, the role of *Bmal1* as a potential modulator of cocaine responses is yet to be addressed. In this context, this work aimed to characterize the role of *Bmal1-KO* in the rewarding and reinforcing effects of cocaine and the molecular alterations linked to this process. Moreover, we investigated the *Bmal1-KO* behavioural phenotype addressing the potential cognitive dysfunctions related to this mutation.

## MATERIALS AND METHODS

### Animals

Heterozygous C57BL/6, (*Bmal1*(+/-)) animals were kindly donated by Stem Cells and Cancer Lab at the IRB (Barcelona) and received at our animal facility, UBIOMEX, PRBB. Animals were placed in pairs in standard cages in a temperature- (21 ± 1°C) and humidity- (55% ± 10%) controlled room subjected to a 12h light/dark cycle, and ad libitum access to food and water. Said animals were used as breeders. Offspring were weaned at postnatal day 21 and housed in groups by sex. After weaning, genotypes were determined; homozygous (*Bmal1*(-/-)), heterozygous (*Bmal1* (+/-)) or WT (*Bmal1* (+/+)). All experiments were carried out in accordance with the guidelines of the European Communities Directive 88/609/EEC regulating animal research. Procedures were approved by the local ethical committee (CEEA-PRBB) and every effort was made to minimize animal suffering and discomfort as well as the number of animals used.

### Drugs

Cocaine was purchased from Alcatel (Madrid, Spain) and was dissolved in sterile physiological saline (0.9%).

### Spontaneous locomotor activity

We used an automatized box to record spontaneous locomotor activity (LE881 IR, Panlab s.l.u., Barcelona, Spain), as previously described^32^. For 30 minutes, mice explored the environment, while two types of movements were registered: ambulations (horizontal), and rearings (vertical). Data were collected in intervals of 5 minutes. Two independent experiments were performed (dark and light phases).

### Y-maze

Spatial working memory of mice was assessed as previously reported^33^. The percentage of alternation was calculated. Two independent experiments were performed (dark and light phases). For detailed description see *Supplementary methods*.

### Novel object recognition (NOR)

The novel object recognition (NOR) task was performed to evaluate short- and long-term memory function (3 and 24 h respectively), as previously described^34^. All sessions were videotaped for subsequent analyses. Videos were assessed with the software BORIS^35^. Two independent experiments were performed (dark and light phases).

### Social interaction test

The test was conducted as previously reported^33,36^. Briefly, the test consisted of three 10-minute phases: habituation, sociability, and social novelty. For sociability, tested mice were introduced to an unfamiliar animal (novel 1) trapped inside one of the holders in one of the chambers. Later, a second intruder mouse (novel 2) was placed in the second empty holder. Time spent in each compartment throughout the sessions was measured. Then, sociability and social novelty scores were calculated. Two independent experiments were performed (dark and light phases).

### Cocaine-induced behavioural sensitization

The procedure consisted of three phases: habituation, acquisition, and challenge ^37^. In the habituation phase, mice were placed individually into actimetry boxes (24×24×24 cm; LE881 IR, Panlab s.l.u., Barcelona, Spain) for 30 min. On the following 5 days (acquisition phase), mice were treated daily with cocaine (7.5 or 10 mg/kg, i.p.) or saline and placed into the actimetry boxes. Spontaneous locomotor activity was recorded for 30 min. A cocaine challenge (7.5 or 10 mg/kg, 30 min) was administered 7 days after the last day of the acquisition phase (also to the saline group). Then, Δ scores were calculated. All mice were tested during the light phase.

### Cocaine-induced conditioned place preference (CPP)

The CPP was performed as previously reported^33,38^. We used two cocaine doses: 5mg/kg and 7.5mg/kg. The CPP score for each animal was then calculated. All mice were tested during the light phase. For detailed description see *Supplementary methods*.

### Cocaine self-administration (SA)

The cocaine SA paradigm was conducted as previously described ^39–41^. Nosepoking into the active hole resulted in a cocaine infusion (0.75 mg/kg/infusion) and nosepoking into the inactive hole had no consequences. All sessions took place during the dark phase. For detailed description see *Supplementary methods*.

### rt-qPCR

Total RNA extraction from medial prefrontal cortex (mPFC) and STR samples was conducted using the trizol method as previously described^40,42^. For the qPCR, we used 20 ng of sample to evaluate the expression of *Clock, Per2, D2R* and *GAPDH* as housekeeping gene. The ΔΔCt was calculated. For detailed description see *Supplementary methods*.

### Western blot (WB)

WBs were performed as described previously^40,41,43^. We evaluated the consequences of *Bmal1-KO* and cocaine administration on the protein expression of GluA1, GluA2, MAO-A, MAO-B, DAT and TH (in the case of STR) in the mPFC and STR. For this, the tissues of animals that underwent the 7.5 mg/kg sensitization were collected after the cocaine challenge. Hence, three treatment conditions were evaluated: basal condition (naïve mice), acute condition (saline during sensitization and cocaine injection during challenge), and chronic condition (cocaine during sensitization and cocaine injection during challenge). Protein expression signals were normalised to the housekeeping control protein in the same samples and expressed in terms of fold-change with respect to control values. Samples were obtained through the light phase (9:00 to 12:00 h). For detailed description see *Supplementary methods*.

## STATISTICAL ANALYSIS

Data were analysed for conditions of normality (Kolmogorov-Smirnov’s test), sphericity (Mauchly’s test) and homoscedasticity (Levene’s test). We analysed the weight results using a three-way ANOVA (*days, sex* and *genotype*). As per the NOR, Y-maze, qPCR and WB, results were analysed using two-tailed unpaired Student’s t-tests. Furthermore, a two-way ANOVA was used to analyse results from the locomotor activity (*time* and *genotype*), sensitization (*compartment* and *genotype*) and CPP (*treatment* and *genotype*). For the cocaine-SA, we analysed the infusions through a two-way ANOVA (*days* and *genotype*) and a three-way ANOVA for the nosepokes (*days, hole*, and *genotype*). Lastly, for the sensitization, the acquisition phase was analysed with a three-way ANOVA (*days, treatment*, and *genotype*) and the challenge, with a two-way ANOVA (*treatment* and *genotype*). When appropriate, repeated measures analyses were applied. If F achieved p < 0.05, the ANOVA was followed by a Bonferroni post-hoc test if the main effect and/or interaction was observed. All possible post-hoc comparisons were evaluated. Statistical analyses were performed using SPSS Statistics v23. Data are expressed as mean ± SEM and a value of p < 0.05 was considered significant.

## RESULTS

### *Bmal1-KO* animals show higher locomotor activity than WT mice during the light phase

We measured locomotor activity in WT and *Bmal1-KO* mice during light and dark phase for 30 minutes. For the horizontal activity (light phase) (Fig. 1A), a two-way ANOVA showed a *time* main effect (F_2.73, 87.4_=12.4, p<0.001) and the interaction *genotype* × *time* (F_5, 160_=4.52, p<0.001). The Bonferroni post-hoc test showed increased activity counts in *Bmal1-KO* group at minute 25 in comparison to WT mice (p<0.05). *Time* main effect (F_3.76, 147_=24.3, p<0.001) and the *genotype* × *time* interaction (F_5, 195_=4.68, p<0.001) were found during the dark phase, but no differences were found between experimental groups. Additional analysis (Student’s t-tests) of the collapsed values accumulated throughout the session showed no differences in horizontal locomotion between experimental groups (Fig. 1B). For the vertical activity (light phase) (Fig. 1C), the two-way ANOVA showed *time*- (F_3.33,106.7_=11.36, p<0.001) and *genotype*- (F_1,32_=7.022, p<0.05) main effects, suggesting elevated vertical movements in the *Bmal1-KO* group. For the dark phase, the two-way ANOVA showed a *time* main effect (F_3.703, 148.1_=20.60, p<0.001). Lastly, the Student’s t-tests of the accumulated vertical values confirmed an increased vertical locomotion in the *Bmal1-KO* mice during the light phase (t_32_=1.108, p<0.05) (Fig. 1D).

**Fig. 1.**
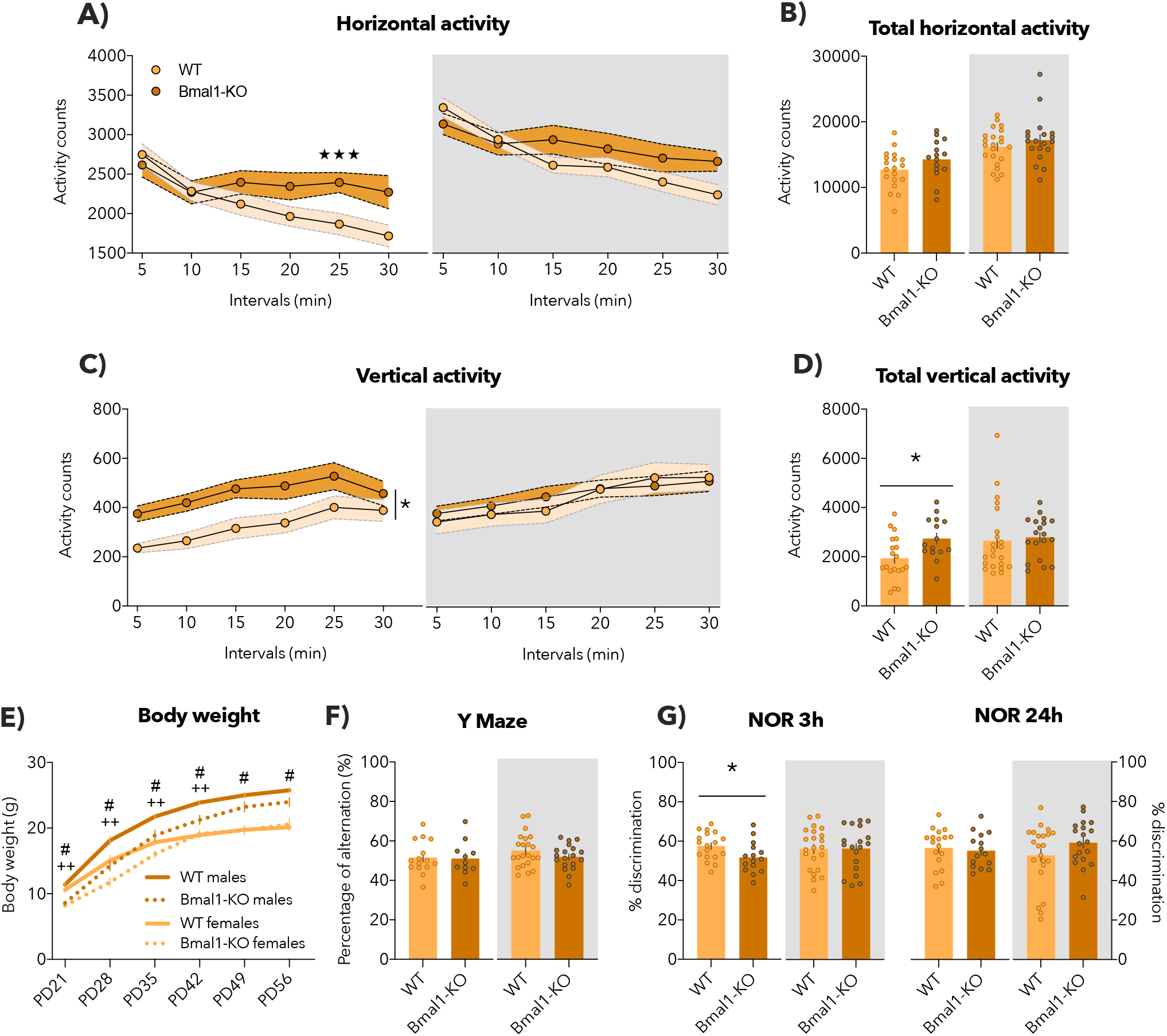
Effects of *Bmal1* gene deletion on locomotor activity, body weight, and memory tasks. Mean of spontaneous locomotor activity (A-D) during the light phase (WT-mice n=19 and *Bmal1-KO*-mice n=15) and the dark phase (WT-mice n=22 and *Bmal1-KO*-mice n=19). *Genotype* main effect (*p< 0.05) and *time* × *genotype* interaction of the ANOVA (★★★ *Bmal1-KO*-males n=8; WT-females n=10; *Bmal1-KO*-females n=5). *Genotype* × *days* interaction of the ANOVA (++ p < 0.01), *days* × *sex* interaction of the ANOVA (# p<0.05). Mean of the percentage of alternation during the Y-maze (F) in the light phase (WT-mice n=19 and *Bmal1-KO*-mice n=15) and dark phase (WT-mice n=22 and *Bmal1-KO*-mice n=19). Percentage of discrimination in the NOR test (G) after 3 h and 24 h (WT-mice n=19 and *Bmal1-KO*-mice n=15 light phase; WT-mice n=22 and *Bmal1-KO*-mice n=19 dark phase) (*p<0.05). Bonferroni post-hoc comparison of the ANOVA. Data are expressed as mean ± SEM. Experiments performed in the dark phase are shaded in the figure.

### *Bmal1-KO* adult mice present unaltered body weight in adulthood

Animals were weighed during 6 specific postnatal days (PD21, PD28, PD35, PD42, PD49 and PD56) to determine the influence of *Bmal1-KO* and sex on body weight (Fig. 1E). The three-way ANOVA yielded the main effects *days* (F_5,145_=923.31, p<0.001), *genotype* (F_1,29_=12.55, p<0.001) and *sex* (F_1,29_=36.04, p<0.001). Also, we found the interactions: *days* × *genotype* (F_5,145_=11.98, p<0.001) and the *days* × *sex* (F_5, 145_=19.22, p<0.001). The post-hoc analysis for the *days* × *genotype* revealed that *Bmal1-KO* mice weighed less than WT animals only during the four first measurements (PD21, PD28, PD35 and PD42) (p<0.01, in all the cases). The *days* × *sex* interaction showed that, as expected, male mice weighed more than females all the days (p<0.05).

### *Bmal1-KO* mice showed normal working memory function

To evaluate working memory, we performed the Y maze (Fig. 1F) during the light and the dark phase. No significant differences were identified in either phase (t_24_=1.476, ns and t_39_=1.359, ns, respectively).

### *Bmal1-KO* animals exhibit short-term memory impairment during the light phase

We performed two measurements for the NOR (Fig. 1G) to evaluate both short- and long-term memory (3 h and 24 h, respectively). The Student’s t-test for the results after 3 h (light phase) revealed that *Bmal1-KO* mice showed reduced ability to discriminate between objects (t_31_=2.227, p<0.05), but yielded no significant differences at 24 h (t31=0.4364, ns). No significative differences were found in the dark phase.

### *Bmal1-KO* mice have reduced social novelty recognition during the light phase

Regarding social behaviour during the light phase, a two-way ANOVA analysis showed normal sociability (*compartment* effect; F_2,96_=174.5, p<0.001) (Fig. 2A), since both groups spent more time in the chamber containing the intruder mouse compared to the two empty chambers. The analysis of the social novelty recognition showed a *compartment* effect (F_2,96_=57.06, p<0.001) and the *compartment* × *genotype* interaction (F_2,96_=7.325, p<0.001) (Fig. 2B). Post-hoc comparisons for the interaction revealed that *Bmal1-KO* did not discriminate between novel 1 and novel 2 mice, while the WT mice did (p<0.001). No significative differences were found in sociability or social novelty recognition during the dark phase.

**Fig. 2.**
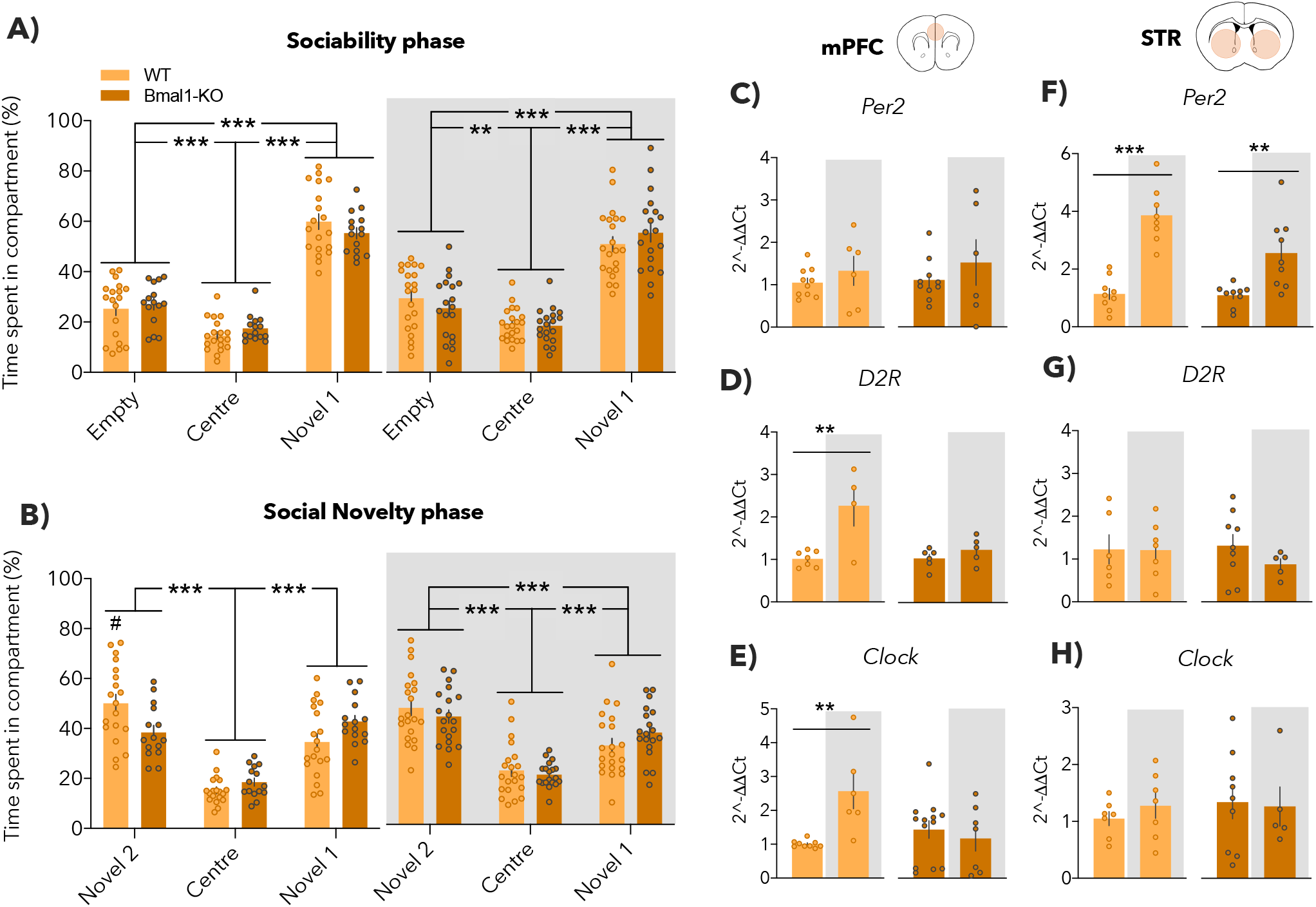
Changes in sociability, social novelty and genes related to circadian rhythms (*Per2, D2R* and *Clock*) on *Bmal1-KO* mice. Time spent in the different compartments in the light phase (WT-mice n=19 and *Bmal1-KO*-mice n=15) and the dark phase (WT-mice n=21 and *Bmal1-KO*-mice n=19). *Compartment* main effect of the ANOVA (**p<0.01; ***p<0.001) and *genotype* × *compartment* interaction of the ANOVA (#p<0.05). Bonferroni post-hoc comparison for the ANOVA. Mean fold change relative to the light condition in each group (C) *Per2*, (D) *D2R* and (E) *Clock* in the mPFC (**p<0.01, Student’s T-test). Mean fold change relative to the light condition in each group (F) *Per2*, (G) *D2R* and (H) *Clock* in the STR (**p<0.01, ***p<0.001, T-test). n=5-15/group, run in duplicate. Data are expressed as mean ± SEM. Experiments performed in the dark phase are shaded in the figure.

### WT mice experience changes in D2R expression throughout the cycle in the mPFC, but these are lost in *Bmal1-KO*

Expression levels of *Per2, D2R*, and *Clock* mRNA at both phases (light and darkness) in the mPFC were examined (Fig. 2C-E). The Student’s T-tests showed elevation of *D2R* (t_9_=3.488, p<0.01) (Fig. 2D) and *Clock* (t_13_=3.759, p<0.01) (Fig. 2E) in the WT group, whereas these circadian rhythms were completely lost in *Bmal1-KO* mice.

### WT and *Bmal1-KO* mice showed increased *Per2* mRNA expression in the STR during the dark phase

We determined the levels of *Per2, D2R*, and *Clock* mRNA in the STR (Fig. 2F-H) during the light- and dark phases. The Student’s t-tests yielded robust *Per2* rhythmicity in the WT (t_15_=7.118, p<0.001) (Fig. 2F) and *Bmal1-KO* mice (t_16_=3.358, p<0.01) (Fig. 2F)

### Cocaine-induced behavioural sensitization in WT and *Bmal1-KO* mice is similar at 7.5 mg/kg dose

The three-way ANOVA for the 7.5 mg/kg dose (Fig. 3A) showed an effect of *day* (F_4,200_=8.679, p<0.001), *treatment* (F_1,50_=80.878, p<0.001) and *days* × *treatment* interaction (F_4,200_=5.791, p<0.001). The post-hoc analysis for the interaction showed that cocaine induced hyperlocomotion every day (p<0.001, in all cases) and higher locomotion on day 5 compared to days 1, 2 and 3 (p<0.05, in all cases). The two-way ANOVA for the challenge (Fig. 3B) showed a cocaine *treatment* effect (F_1,50_=12.54, p<0.001). Additionally, we compared the results of day 1, day 5 and challenge (Fig. 3C) with a two-way ANOVA. The analysis showed effect of *days* (F_2,100_=40.933, p<0.001), *treatment* (F_1,50_=66.817, p<0.001) and the *days* × *treatment* interaction (F_2,100_=7.585, p<0.001). Post-hoc comparisons for the interaction showed hyperlocomotion in cocaine-treated animals all days (p<0.001, in all cases), higher locomotion on day 5 compared to day 1 (p<0.001), and increased hyperlocomotion on challenge day compared to day 1 (p<0.001).

**Fig. 3.**
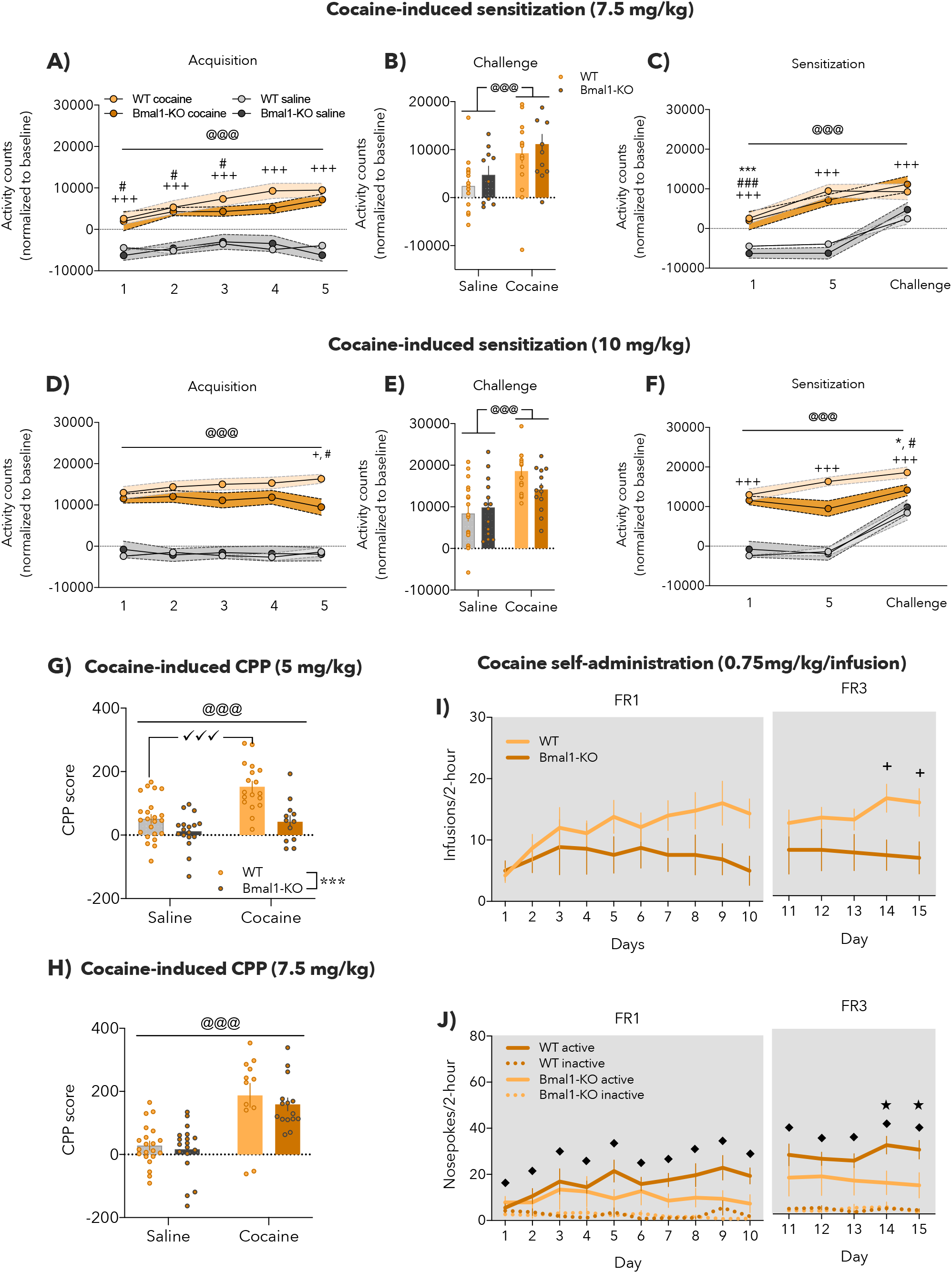
*Bmal1-KO* induced alterations in the rewarding effects of cocaine in different paradigms. *Sensitization*. Locomotor activity (n=10-18/group) during behavioral sensitization sessions (A,D; 7.5 and 10 mg/kg, respectively), after the cocaine challenge (B,E; 7.5 and 10 mg/kg, respectively) and the results of day 1, day 5 and the challenge day (C,F; 7.5 and 10 mg/kg, respectively). *Sensitization 7*.*5 mg/kg*. (A) *Treatment* effect (@@@p<0.001), *days* × *treatment* (cocaine vs saline, +++p<0.001; difference from day 5, #p<0.05). (B) *Treatment* effect (@@@p<0.001). (C) *Treatment* effect (@@@p<0.001), *days* × *treatment* (cocaine vs saline, +++p<0.001; difference from day 5, ###p<0.001; day 1 vs. challenge, ***p<0.001). *Sensitization 10 mg/kg*. (D) *Treatment* effect (@@@p<0.001), *days* × *genotype* (day 5 vs. day 1 on WT-mice, +p<0.05; WT vs KO, #p<0.05). (E) *Treatment* effect (@@@p<0.001). (F) *Treatment* effect (@@@p<0.001), *days* × treatment (cocaine vs saline, +++p<0.001; day 1 vs. challenge, *p<0.05; day 5 vs. challenge, #p<0.05). *Cocaine-induced conditioned place preference*. CPP is expressed as differences of time spent in the saline/cocaine-paired compartment between the test day and the pre-test day at (G) 5 mg/kg and (H) 7.5 mg/kg of cocaine. *Treatment* (@@@p<0.001) and *genotype* (***<0.001) main effect and *treatment* × *genotype* interaction of the ANOVA (✓✓✓ p<0.001) (n=12-23/group). *Cocaine self-administration*. Mean of (I) infusions during the FR1 and FR3 and mean of (J) nosepokes on the FR1 and along the 10-day of self-adminsitration (WT-mice n=9, *Bmal1-KO*-mice n=7). *Days* × *genotype* (+, p<0.05), *days* × *hole* (♦, p<0.05) and *days* × *genotype* × *hole* (★p< 0.05). Bonferroni post-hoc comparison for the ANOVA. Data are expressed as mean ± SEM. Experiments performed in the dark phase are shaded in the figure.

### *Bmal1-KO* mice show reduced cocaine-induced sensitization at a dose of 10 mg/kg

The three-way ANOVA revealed a *treatment* effect (F_1,58_=142.477, p<0.001) and *days* × *genotype* interaction (F_4,232_=3.579, p<0.01) (Fig. 3D). The Bonferroni’s test for the interaction showed that cocaine induced higher locomotor activity on day 5 compared to day 1 only in WT (p<0.05) while KO animals showed no differences in locomotion along the days. Additionally, the post-hoc test yielded a higher cocaine-induced hyperlocomotion in WT mice on day 5 compared to KO (p<0.05). The two-way ANOVA for the challenge (Fig. 3E) showed a cocaine *treatment* effect (F_1,58_=18.80, p<0.001).

The two-way ANOVA for results obtained on day 1, day 5 and challenge (Fig. 3F) revealed an effect of *days* (F_2,116_=53.283, p<0.001), *treatment* (F_1,58_=86.219, p<0.001) and the *days* × *treatment* interaction (F_2,116_=14.698, p<0.001). The Bonferroni’s test for the interaction showed higher locomotion in cocaine-treated animals every day (p<0.001, in all cases), and increased locomotor activity during challenge compared to day 1 (p<0.05) and 5 (p<0.05).

### *Bmal1-KO* mice exhibit reduced rewarding effects of cocaine at low doses (5 mg/kg)

We evaluated the cocaine CPP at two different doses: 5 mg/kg (Fig. 3G) and 7.5 mg/kg (Fig. 3H) during the light phase. The two-way ANOVA for the 5 mg/kg dose showed a significant effect of *treatment* (F_1,66_=16.15, p<0.01), *genotype* (F_1,66_=20.93, p<0.01), and interaction between these factors (F_1,66_=4.755, p<0.05). The Bonferroni’s test revealed that WT animals showed increased preference towards the cocaine-paired compartment (p<0.001) while KO mice did not develop cocaine CPP. The two-way ANOVA for 7.5 mg/kg revealed a *treatment* effect (F_1,62_=47.12, p<0.001), meaning that both groups increased the time spent in the cocaine-paired compartment similarly.

### *Bmal1-KO* mice display reduced cocaine-seeking behaviour in the self-administration paradigm

WT and *Bmal1-KO* mice were trained to self-administer 0.75 mg/kg/inf cocaine for 10 *days* under a fixed ratio (FR) 1 reinforcement schedule during the dark phase. Afterwards, animals underwent FR3 for 5 days.

The two-way ANOVA for infusions in the FR1 (Fig. 3I) revealed a *day* effect (F_9,126_=2.700, p<0.01), meaning that animals increased the number of infusions throughout the days. The two-way ANOVA for the FR3 (Fig. 3I) revealed a *day* × *genotype* interaction (F_4,56_=3.097, p<0.05). The Bonferroni’s test showed higher number of infusions made by WT on day 14 and 15 (p<0.05, in all cases).

The three-way ANOVA for nosepokes during FR1 (Fig. 3J) indicated a *hole* effect (F_1,14_=29.412, p<0.001) and the *day* × *hole* interaction (F_9,126_=2.203, p<0.05). Post-hoc comparisons confirmed that animals discriminated between the active and the inactive holes from day 1 until the end of the FR1 phase. The three-way ANOVA for the FR3 (Fig. 3J) yielded a *hole* effect (F_1,14_=36.663, p<0.001), the *day* × *genotype* interaction (F_4,56_=4.178, p<0.01) and the *day* × *genotype* × *hole* interaction (F_4,56_=2.526, p<0.05). The post-hoc test for the triple interaction yielded that WT performed more active nosepokes than KO on day 14 and 15 (p<0.05, in all cases). Additionally, the post-hoc test indicated that both groups discriminated between holes from the beginning until the end of the FR3 cocaine-self-administration (p<0.05, in all cases).

### *Bmal1-KO* showed reduced GluA1 (mPFC) protein level after chronic cocaine administration (7.5 mg/kg)

The Student’s T-tests for the mPFC revealed decreased GluA1 protein expression (t_9_=3.312, p<0.01) (Fig. 4A) after chronic cocaine administration in the *Bmal1-KO* mice compared to WT.

**Fig. 4.**
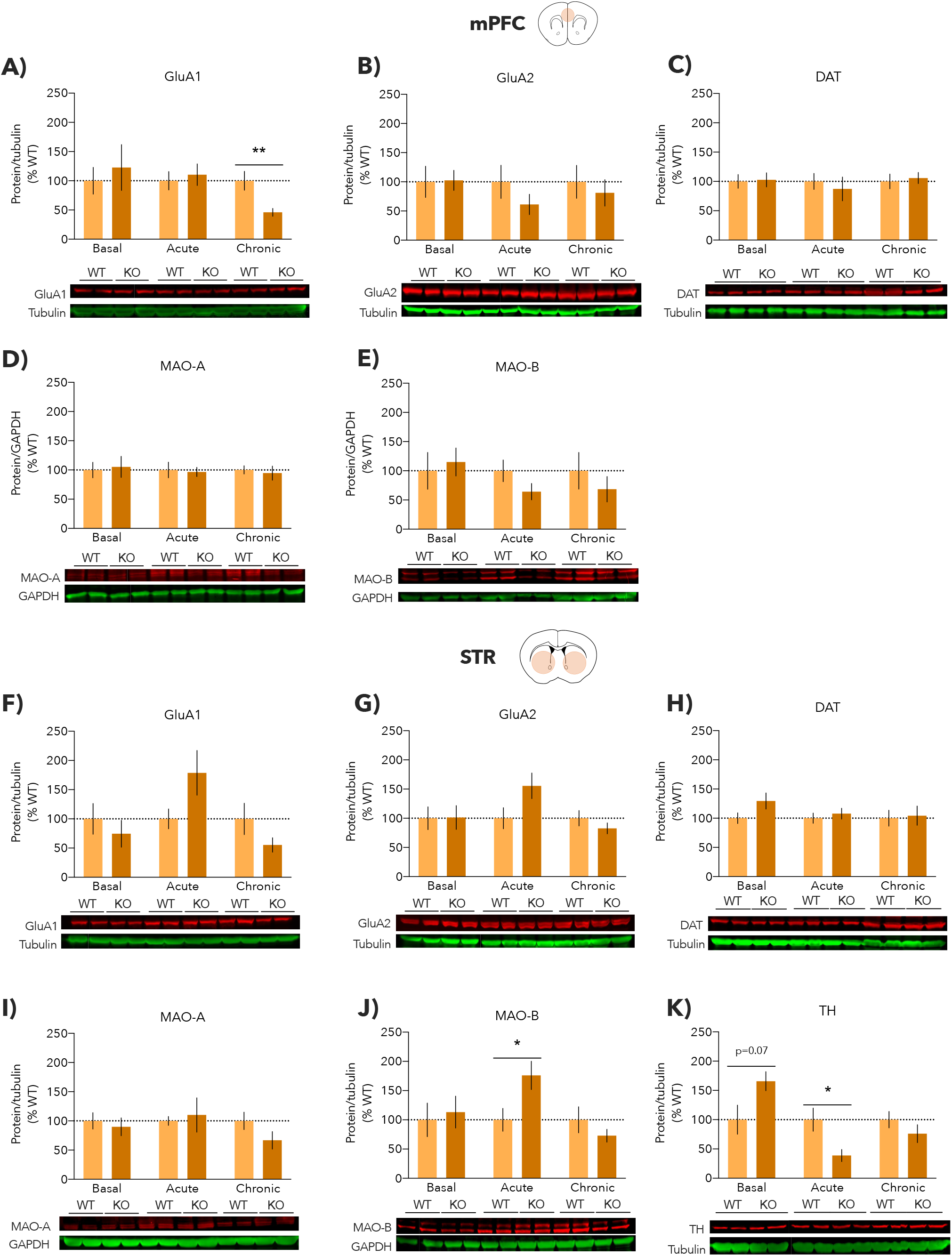
Western blot analyses in the medial prefrontal cortex and striatum of WT mice and *Bmal1-KO* animals under basal conditions and after acute or repeated cocaine administration. We obtained the mPFC and the STR after the cocaine challenge (7.5 mg/kg sensitization). Three treatment conditions: basal condition (naïve mice), acute condition (saline during the sensitization and cocaine injection for the challenge), and chronic condition (cocaine during the sensitization and cocaine injection for the challenge). Mean fold change relative to the WT group of (A) GluA1, (B) GluA2, (C) DAT, (D) MAO-A, (E) MAO-B in the mPFC. Mean fold change relative to the WT group (F) GluA1, (G) GluA2, (H) DAT, (I) MAO-A, (J) MAO-B, (K) TH in the STR. Student’s T-test comparing WT and *Bmal1-KO* mice in each treatment condition (*p<0.05, **p<0.01). The protein of interest in red and housekeeping protein in green. Bonferroni post-hoc comparison for the ANOVA. Data are expressed as mean ± SEM (n = 4-6, run in duplicate or triplicate).

### *Bmal1-KO* showed reduced TH and increased MAO-B in STR after a single i.p. cocaine injection (7.5 mg/kg)

The Student’s T-test analyses for the STR yielded that acute cocaine administration increased protein expression of MAO-B (t_9_=2.483, p<0.05) (Fig. 4J) and reduced TH (Fig. 4K) protein level (t_10_=2.747, p<0.05) in *Bmal1-KO* mice in comparison with WT.

## DISCUSSION

Our results revealed that *Bmal1-KO* mice show less cocaine sensitization at 10 mg/kg, impaired cocaine CPP at 5 mg/kg and reduced cocaine-seeking behaviour in the self-administration paradigm. Moreover, these animals had increased locomotor activity during the light phase, reduced short-term memory and diminished novel social interaction. Regarding molecular results, *Bmal1-KO* mice display lost circadian rhythmicity in *D2R* and *Clock* mRNA expression in the mPFC. Moreover, acute cocaine administration induced enhanced protein expression of MAO-B (STR) and reduced the synthesis of TH (mPFC) in *Bmal1-KO*. Lastly, chronic cocaine administration decreased protein expression of GluA1 (mPFC) in *Bmal1-KO* in comparison with WT.

Generally, we found the biggest behavioural differences during the light phase. This is in line with a previous study reporting that clock rhythmicity is maintained during the dark phase in the absence of *Bmal1* but not during the light phase of the cycle ^44^. Circadian clock genes can have an impact on behaviours like locomotion, cognition, and mood ^12,22–24,45^. Regarding locomotion, other authors using full-body *Bmal1-KO* animals reported reduced locomotor activity compared to WT ^2,12,44^ when locomotion activity was recorded 24 h in their home cage. In contrast, other authors using different mouse models of circadian gene deletion which assess locomotion at specific time points, instead of the entire cycle (24 h), revealed: slow locomotor habituation to a novel environment in mutant mice compared to WT ^23^, less travelled distance ^23^ and hyperactivity in rearing and locomotor activity in a novel environment ^30,45^. In line with these reports, we observed increased spontaneous locomotor activity in *Bmal1-KO* mice compared to WT animals, confirming their reduced ability to habituate to novel environments.

As mentioned before, *Bmal1* gene is involved in cognition. Here, we reported impaired NOR after 3 h and failed capacity to discern between the familiar and novel mouse during the social novelty test. *Bmal1-KO* accelerates ageing, and this biological process accompanies memory decline. Indeed, Kondratova *et al*. (2010) described alterations in short-term memory in *Bmal1-KO* mice^45^, and a forebrain-specific *Bmal1-KO* showed deficient acquisition in Barnes maze and novel object location memory^46^. Then, our results are in line with memory decline and reduced ability to form new short-time memories observed in previous studies.

Concerning cocaine results, when we used a dose of 7.5 mg/kg for the cocaine-induced sensitization, both WT and *Bmal1-KO* mice showed locomotor sensitization to the drug. Nevertheless, when a higher dose was administered (10 mg/kg), *Bmal1-KO* cocaine-induced sensitization was impaired. In fact, they showed reduced locomotor activity on day 5 compared to WT mice. Additionally, WT mice displayed a higher locomotor sensitization than the mutant group independently of the dose. The initiation of cocaine sensitization involves alterations in dopamine transmission in the ventral tegmental area, STR and mPFC^47^. Dopamine biosynthesis is negatively regulated by circadian nuclear receptor REV-ERBα^23^ and CLOCK^17,30^ through transcriptional regulation of *TH* (the rate-limiting enzyme in dopamine synthesis). *Rev-erbα* expression is controlled by the binding of clock proteins complex (BMAL1:CLOCK or BMAL1:NPAS2)^22^ to their E-box elements. Therefore, it is expected that in absence of *Bmal1*, animals will exhibit increased TH and dopamine levels. Accordingly, we observed a statistical tendency to an increase in TH protein basal expression in the STR of *Bmal1-KO*.

Additionally, other authors observed that TH inhibition leads to the rescue of some dopamine-induced alterations in *Per2* mutants’ mice^24^. Here, we reported reduced TH protein expression in the STR of *Bmal1-KO* mice after acute cocaine administration (but not after chronic administration). We suggest a TH reduction in *Bmal1-KO* mice as a compensatory mechanism to counter the cocaine-induced dopamine activation. However, this will be only temporary because after repeated cocaine administration, *Bmal1-KO* cannot afford this compensation.

On the other hand, behavioural sensitization to cocaine also requires a decrease in drug-induced cortical dopamine concentrations^47^. The MAO is an enzyme that plays a key role in the degradation of dopamine ^48^. There are two MAO <u>isoenzymes</u>: MAO-A and MAO-B^48^. Although both enzymes catalyse most of the biogenic amines, the contribution of MAO-B to dopamine degradation is controversial^49^. It is believed that MAO-A, but not MAO-B, mainly contributes to striatal dopamine breakdown^49^. Besides, *Bmal1* together with NPAS2 positively regulates MAO-A transcription^22^, but no MAO-B’s^24^. Here, we observed that after acute cocaine administration, *Bmal1-KO* mice showed a strong MAO-B elevation in the STR in comparison with WT. Therefore, there is a possibility that in the absence of *Bmal1*, MAO-B attempts to catalyse the dopamine elevation together with the reduction of TH expression.

The reduction of dopamine degradation suggested in the present and previous studies ^24^ could result in a rise of the dopamine levels in brain areas such as mPFC and STR. Therefore, higher cocaine doses could result aversive in *Bmal1-KO* mice. This could explain why *Bmal1-KO* mice showed reduced cocaine-induced locomotor sensitization at the highest cocaine dose (10 mg/kg), including reduced locomotion on day 5 compared to day 1.

Furthermore, other studies reported that D2R in the mPFC have a key role in cocaine-induced responses^47^. Beyer and Stekettee (2002) demonstrated that intra-mPFC injection of a D2R agonist blocked the initiation and attenuated the expression of locomotor sensitization to cocaine^47^. Moreover, diurnal variations in DAT and D2R are critical for dopamine activity and reward regulation 50. Dopamine receptors expression is regulated by NPAS251 which acts as a CLOCK paralog52. Here, we observed altered circadian rhythmicity in *D2R* and *Clock* in the mPFC of *Bmal1-KO*. Those alterations could be adding to the impaired cocaine-related responses.

Because of the above-mentioned alterations in dopamine system components observed in these mutant mice, we explored the role of *Bmal1* in the rewarding effects of cocaine by using two different behavioural approaches: CPP and SA.

Our results showed no CPP at the lowest cocaine dose (5 mg/kg) and reduced cocaine seeking during the SA paradigm in the *Bmal1-KO* mice. These results suggest reduced cocaine’s motivational effects in these mutant mice. Another group evaluating cocaine-induced adaptations in circadian genes reported that increased cocaine consumption in the SA paradigm is accompanied by elevated expression of several circadian genes like *Bmal1* in the STR of rats29. Hence, it is not surprising to find a reduced cocaine SA in *Bmal1* KO mice. Other authors reported decreased GluA1 in the STR of *Clock* mutants and they suggest these post-synaptic changes as secondary adaptations to the increased dopaminergic signalling53.

Finally, GluA1 is closely related to cocaine sensitization and increased drug reward40,43. Here, we also reported that after chronic cocaine administration, *Bmal1-KO* mice showed reduced GluA1 in the mPFC. This is consistent with the reduced cocaine rewarding effects and the altered cocaine response observed in the paradigms that we explored.

## CONCLUSIONS

Taken together, our results confirm the key role of *Bmal1* as a molecular regulators of the brain’s reward circuitry and/or circadian rhythmicity. Moreover, we demonstrated a key role of *Bmal1* in the motivational effects of cocaine since *Bmal1-KO* mice display blunted CPP at 5 mg/kg and decreased cocaine seeking in the SA paradigm. We suggest that reduced cocaine’s rewarding effects in these mutant mice might be related to *Bmal1* role as an expression regulator of MAO and TH, two essential enzymes involved in dopamine metabolism.

## ABBREVIATIONS

SCN: suprachiasmatic nucleus
KO: Knockout
WT: wild type
TH: tyrosine hydroxylase
MAO-A: monoamine oxidase A
MAO-B: monoamine oxidase B
D2R: dopamine receptors 2
D3R: dopamine receptors 3
STR: striatum
NPAS2: neuronal PAS domain protein 2
CPP: conditioned place preference
NOR: novel object recognition
SA: self-administration
mPFC: medial prefrontal cortex
GluA1: AMPA receptor subunit 1
GluR2: AMPA receptor subunit 2
DAT: dopamine transporter
PD: postnatal day

## ACKNOWLEDGEMENTS

This study was supported by the Ministerio de Economia y Competitividad (grant number PID2019-104077RB-100), Ministerio de Sanidad (Retic-ISCIII, grant number RD16/017/010; RICORS, grant number RD21/0009/0001, and Plan Nacional sobre Drogas 2018/007). L.A-Z and I.G.-L. received a FPI grants (BES-2017-080066 and PRE2020-091923) from the Ministerio de Economia y Competividad. P.S.W. was supported by grant RYC2019-026661-I funded by MCIN/AEI/10.13039/501100011033 and by “ESF Investing in your future”. The Department of Medicine and Health Sciences (MELIS-UPF) is a “Unidad de Excelencia María de Maeztu” funded by the AEI (CEX2018-000792-M). O.V. is recipient of an ICREA Academia Award (Institució Catalana de Recerca i Estudis Avançats, Generalitat de Catalunya).

## AUTHOR CONTRIBUTIONS

A.C.Z., L.A.Z., L.C., P.S.W., S.A.B. and O.V. were responsible for the study concept and design. A.C.Z., L.A.Z., L.C., I.G.L. carried out the experimental studies. A.C.Z, L.A.Z, I. G. L and O.V. drafted the manuscript and participated in the interpretation of findings. All authors critically reviewed the content and approved the final version for publication.

## CONFLICT OF INTEREST

The authors declare no conflicts of interest.

## Notes

### Competing Interest Statement

The authors have declared no competing interest.

## REFERENCES

1. Hastings, M. H., Maywood, E. S. & Brancaccio, M. Generation of circadian rhythms in the suprachiasmatic nucleus. Nature Reviews Neuroscience vol. 19 453–469 (2018).

2. Welz, P. S. et al. BMAL1-Driven Tissue Clocks Respond Independently to Light to Maintain Homeostasis. Cell 177, 1436-1447.e12 (2019).

3. Kondratov, R. V., Shamanna, R. K., Kondratova, A. A., Gorbacheva, V. Y. & Antoch, M. P. Dual role of the CLOCK/BMAL1 circadian complex in transcriptional regulation. The FASEB Journal 20, 530–532 (2006).

4. Reppert, S. M. & Weaver, D. R. Coordination of circadian timing in mammals. Nature 418, 935–941 (2002).

5. Patke, A., Young, M. W. & Axelrod, S. Molecular mechanisms and physiological importance of circadian rhythms. Nature Reviews Molecular Cell Biology (2019) doi:10.1038/s41580-019-0179-2.

6. Preitner, N. et al. The orphan nuclear receptor REV-ERBalpha controls circadian transcription within the positive limb of the mammalian circadian oscillator. Cell 110, 251–260 (2002).

7. Sato, T. K. et al. A functional genomics strategy reveals Rora as a component of the mammalian circadian clock. Neuron 43, 527–537 (2004).

8. Liu, A. C. et al. Intercellular Coupling Confers Robustness against Mutations in the SCN Circadian Clock Network. Cell 129, 605–616 (2007).

9. Bunger, M. K. et al. Mop3 Is an Essential Component of the Master Circadian Pacemaker in Mammals. Cell 103, 1009–1017 (2000).

10. Kondratov, R. V, Kondratova, A. A., Gorbacheva, V. Y., Vykhovanets, O. V & Antoch, M. P. Early aging and age-related pathologies in mice deficient in BMAL1, the core componentof the circadian clock. Genes Dev 20, 1868–73 (2006).

11. Bunger, M. K. et al. Progressive arthropathy in mice with a targeted disruption of the Mop3/Bmal-1 locus. genesis 41, 122–132 (2005).

12. Haque, S. N., Booreddy, S. R. & Welsh, D. K. Effects of BMAL1 Manipulation on the Brain’s Master Circadian Clock and Behavior. Yale J Biol Med 92, 251–258 (2019).

13. Charrier, A., Olliac, B., Roubertoux, P. & Tordjman, S. Clock Genes and Altered Sleep-Wake Rhythms: Their Role in the Development of Psychiatric Disorders. Int J Mol Sci 18, (2017).

14. DePoy, L. M., McClung, C. A. & Logan, R. W. Neural Mechanisms of Circadian Regulation of Natural and Drug Reward. Neural Plast 2017, 5720842 (2017).

15. Falcón, E. & McClung, C. A. A role for the circadian genes in drug addiction. Neuropharmacology vol. 56 91–96 (2009).

16. Freyberg, Z. & Logan, R. W. The intertwined roles of circadian rhythmsand neuronal metabolism fueling drug reward and addiction. Current Opinion in Physiology 5, 80–89 (2018).

17. Parekh, P. K., Ozburn, A. R. & McClung, C. A. Circadian clock genes: effects on dopamine, reward and addiction. Alcohol 49, 341–9 (2015).

18. Gordon, H. W. Differential Effects of Addictive Drugs on Sleep and Sleep Stages. J Addict Res (OPAST Group) 3, (2019).

19. McClung, C. & Becker-Krail, D. Implications of circadian rhythm and stress in addiction vulnerability. F1000Res 5, (2016).

20. Sleipness, E. P., Jansen, H. T., Schenk, J. O. & Sorg, B. A. Time-of-day differences in dopamine clearance in the rat medial prefrontal cortex and nucleus accumbens. Synapse 62, 877–885 (2008).

21. Castañeda, T. R., Marquez De Prado, B., Prieto, D. & Mora, F. Circadian rhythms of dopamine, glutamate and GABA in the striatum and nucleus accumbens of the awake rat: modulation by light. J Pineal Res 36, 177–185 (2004).

22. Albrecht, U. Molecular mechanisms in mood regulation involving the circadian clock. Frontiers in Neurology vol. 8 (2017).

23. Chung, S. et al. Impact of circadian nuclear receptor REV-ERBα on midbrain dopamine production and mood regulation. Cell 157, 858–868 (2014).

24. Hampp, G. et al. Regulation of Monoamine Oxidase A by Circadian-Clock Components Implies Clock Influence on Mood. Current Biology 18, 678–683 (2008).

25. Akhisaroglu, M., Kurtuncu, M., Manev, H. & Uz, T. Diurnal rhythms in quinpirole-induced locomotor behaviors and striatal D2/D3 receptor levels in mice. Pharmacology Biochemistry and Behavior 80, 371–377 (2005).

26. Ikeda, E. et al. Molecular Mechanism Regulating 24-Hour Rhythm of Dopamine D3 Receptor Expression in Mouse Ventral Striatum. Molecular Pharmacology 83, 959–967 (2013).

27. Falcon, E., Ozburn, A., Mukherjee, S., Roybal, K. & McClung, C. A. Differential regulation of the period genes in striatal regions following cocaine exposure. PLoS One 8, e66438 (2013).

28. Wang, D.-Q. et al. Effects of chronic cocaine exposure on the circadian rhythmic expression of the clock genes in reward-related brain areas in rats. Behavioural Brain Research 363, 61–69 (2019).

29. Lynch, W. J., Girgenti, M. J., Breslin, F. J., Newton, S. S. & Taylor, J. R. Gene profiling the response to repeated cocaine self-administration in dorsal striatum: A focus on circadian genes. Brain Research 1213, 166–177 (2008).

30. McClung, C. A. et al. Regulation of dopaminergic transmission and cocaine reward by the Clock gene. Proc Natl Acad Sci U S A 102, 9377–9381 (2005).

31. DePoy, L. M. et al. Circadian-Dependent and Sex-Dependent Increases in Intravenous Cocaine Self-Administration in Npas2 Mutant Mice. J Neurosci 41, 1046–1058 (2021).

32. Gracia-Rubio, I. et al. Maternal Separation Impairs Cocaine-Induced Behavioural Sensitization in Adolescent Mice. PLoS One 11, e0167483 (2016).

33. Alegre-Zurano, L., Martín-Sánchez, A. & Valverde, O. Behavioural and molecular effects of cannabidiolic acid in mice. Life Sci 259, (2020).

34. García-Baos, A., Puig-Reyne, X., García-Algar, Ó. & Valverde, O. Cannabidiol attenuates cognitive deficits and neuroinflammation induced by early alcohol exposure in a mice model. Biomedicine and Pharmacotherapy 141, (2021).

35. Friard, O. & Gamba, M. BORIS: a free, versatile open-source event-logging software for video/audio coding and live observations. Methods in Ecology and Evolution 7, 1325–1330 (2016).

36. Portero-Tresserra, M. et al. Maternal separation increases alcohol-drinking behaviour and reduces endocannabinoid levels in the mouse striatum and prefrontal cortex. European Neuropsychopharmacology 28, 499–512 (2018).

37. Cantacorps, L., Montagud-Romero, S., Luján, M. Á. & Valverde, O. Prenatal and postnatal alcohol exposure increases vulnerability to cocaine addiction in adult mice. Br J Pharmacol 177, 1090–1105 (2020).

38. Luján, M. Á. M. Á., Castro-Zavala, A., Alegre-Zurano, L. & Valverde, O. Repeated Cannabidiol treatment reduces cocaine intake and modulates neural proliferation and CB1R expression in the mouse hippocampus. Neuropharmacology 143, 163–175 (2018).

39. Alegre-Zurano, L. et al. Cannabidiol decreases motivation for cocaine in a behavioral economics paradigm but does not prevent incubation of craving in mice. Biomedicine & Pharmacotherapy 148, 112708 (2022).

40. Castro-Zavala, A., Martín-Sánchez, A., Montalvo-Martínez, L., Camacho-Morales, A. & Valverde, O. Cocaine-seeking behaviour is differentially expressed in male and female mice exposed to maternal separation and is associated with alterations in AMPA receptors subunits in the medial prefrontal cortex. Progress in Neuro-Psychopharmacology and Biological Psychiatry 109, (2021).

41. Castro-Zavala, A., Martín-Sánchez, A. & Valverde, O. Sex differences in the vulnerability to cocaine’s addictive effects after early-life stress in mice. Eur Neuropsychopharmacol 32, 12–24 (2020).

42. Cardenas-Perez, R. E. et al. Maternal overnutrition by hypercaloric diets programs hypothalamic mitochondrial fusion and metabolic dysfunction in rat male offspring. Nutr Metab (Lond) 15, 38 (2018).

43. Castro-Zavala, A., Martín-Sánchez, A., Luján, M. Á. & Valverde, O. Maternal separation increases cocaine intake through a mechanism involving plasticity in glutamate signalling. Addiction biology 26, e12911 (2021).

44. Husse, J., Leliavski, A., Tsang, A. H., Oster, H. & Eichele, G. The light-dark cycle controls peripheral rhythmicity in mice with a genetically ablated suprachiasmatic nucleus clock. FASEB Journal 28, 4950–4960 (2014).

45. Kondratova, A. A., Dubrovsky, Y. v., Antoch, M. P. & Kondratov, R. v. Circadian clock proteins control adaptation to novel environment and memory formation. Aging 2, 285–297 (2010).

46. Snider, K. H. et al. Modulation of learning and memory by the targeted deletion of the circadian clock gene Bmal1 in forebrain circuits. Behavioural Brain Research 308, 222–235 (2016).

47. Beyer, C. E. & Steketee, J. D. Cocaine sensitization: Modulation by dopamine D2 receptors. Cerebral Cortex 12, 526–535 (2002).

48. Mattevi, A. Monoamine Oxidase. in Encyclopedia of Biological Chemistry: Second Edition 185–187 (Elsevier, 2013). doi:10.1016/B978-0-12-378630-2.00375-3.

49. Cho, H.-U. et al. Redefining differential roles of MAO-A in dopamine degradation and MAO-B in tonic GABA synthesis. Exp Mol Med 53, 1148–1158 (2021).

50. Ferris, M. J. et al. Dopamine transporters govern diurnal variation in extracellular dopamine tone. Proc Natl Acad Sci U S A 111, E2751–9 (2014).

51. Ozburn, A. R. et al. Direct regulation of diurnal Drd3 expression and cocaine reward by NPAS2. Biol Psychiatry 77, 425–433 (2015).

52. DeBruyne, J. P., Weaver, D. R. & Reppert, S. M. CLOCK and NPAS2 have overlapping roles in the suprachiasmatic circadian clock. Nature Neuroscience 10, 543–545 (2007).

53. Dzirasa, K. et al. Lithium ameliorates nucleus accumbens phase-signaling dysfunction in a genetic mouse model of mania. J Neurosci 30, 16314–23 (2010).

