## Supplementary methods for "*Bmal1*-knockout mice exhibit reduced cocaine-seeking behaviour and cognitive impairments"

**Y-maze**

The task was performed by introducing an animal in the center of a Y-shaped maze with three equal arms, each 39.5 cm long and separated by 120º angles, and were allowed to freely explore it for 8 minutes. Alternation was described as 3 consecutive entries in the 3 different arms. Then, the percentage of alternation was calculated. Two independent experiments were performed (dark and light phases).

**Cocaine-induced conditioned place preference (CPP)**

The apparatus used for this purpose consisted of two compartments (30x29x35cm each) with different visual and tactile properties (Cibertec S.A., Madrid, Spain). One chamber had white walls with textured flooring consisting of prism shapes, whereas the other one, had black walls and a smooth floor. Both were equipped with infrared detectors that allowed the location of the animals throughout the procedure. Mice were placed in the central compartment and had free access to both compartments of the apparatus. During the conditioning phases, mice were randomly assigned to one of the experimental groups, saline (vehicle) or cocaine (5mg/kg or 7.5mg/kg). Mice received an i.p. injection of cocaine (4 cocaine pairings, 8 days) immediately before being placed into one of the two conditioning compartments for 20 min. Mice were given a saline injection on alternate days and placed in the other container for 20 min. Control animals received saline every day. The CPP score for each animal was then calculated. All mice were tested during the light phase.

**Cocaine self-administration (SA)**

*Apparatus for self-administration experiments.* The self-administration was carried out in mouse operant chambers containing two holes; one was defined as active and the other as inactive. Nosepoking into the active hole produced a cocaine infusion that was paired with with a light placed above the active hole. Nosepoking into the inactive hole had no consequences. The side on which the active/inactive hole was placed was counterbalanced.

*Cocaine self-administration procedure.* At least 3 days after surgery, animals were trained, on a fixed ratio 1, to self-administer cocaine (0.75 mg/kg/infusion) during 10 daily sessions (2 h). Cocaine infusion was delivered in 20 µL over 2s via a syringe located on a microinfusion pump (PHM-100A, Med-Associates, Georgia, VT, USA) and connected via Tygon tubing (0.96 mm outer diameter, Portex Fine Bore Polythene Tubing, Portex Limited, Kent, England) to a liquid swivel (375/25, Instech Laboratories, Plymouth Meeting, PA, USA) and the mouse intravenous catheter. In order to avoid overdoses, mice will receive a maximum of 150 infusions and each reinforcement was followed by a 15 s time-out period, in which no cocaine infusions were delivered. The session started with a cocaine priming injection and a 4 s presentation of the light cue, situated above the active hole. All sessions took place during the dark phase.

**RNA extraction, cDNA sequencing and rt-qPCR**

Total RNA extraction from medial prefrontal cortex (mPFC) and STR samples was conducted using the trizol method as previously described45,47. rt-PCR was performed by High-Capacity cDNA Reverse Transcription Kit (Applied Biosystems, Foster City, CA) using random primers and following standardized protocols. As per the qPCR, 20 ng of sample were loaded in addition to the following reagents: LightCycler SBYR green 480 Master Mix (Roche LifeScience, Product No. 04707516001) and the specific primers for Clock, Per2, D2R and GAPDH as housekeeping gene (Integrated DNA Technologies, Inc.) (Table S1). The qPCR was performed in LightCycler® 480 Instrument II (Roche LifeScience). Then, the ΔΔCt was calculated.

**Western blot**

We sought to identify the consequences of Bmal1-KO and cocaine administration on the protein expression of GluA1, GluA2, MAO-A, MAO-B, DAT and TH (in the case of STR) in the mPFC and STR. We obtained the tissues of the animals that underwent the 7.5 mg/kg sensitization, after the cocaine challenge. Hence, three treatment conditions were evaluated: basal condition (naïve mice), acute condition (saline during the sensitization and cocaine injection for the challenge), and chronic condition (cocaine during the sensitization and cocaine injection for the challenge). Western blots were performed as described previously45,46,48. Samples were homogenized in cold lysis buffer and protein samples (16 µg) were mixed with 5X loading buffer, loaded and run on SDS-PAGE 10% and transferred to PVDF membranes. Membranes were blocked with BSA 5% for 1 h at room temperature and incubated overnight at 4ºC with primary antibodies (Table S2). Then, membranes were incubated for 1 h with their respective secondary fluorescent antibodies (Table S3). Protein expression was registered using an Amersham™ Typhoon™ scanner and quantified using Image Studio Lite software v5.2 (LICOR, USA). Protein expression signals were normalised to the detection of housekeeping control protein (Tubulin or GAPDH) in the same samples and expressed in terms of fold-change with respect to control values.

**Table S1**

| Gene | Primer sequence (5`→3’) |
|---|---|
| Clock forward | GGTCAAGGGCTACAGATGTTT |
| Clock reverse | CAGGTGTGAGTTGCTGGATATTA |
| Per2 forward | CAACAACCCACACACCAAAC |
| Per2 reverse | CTCGATCAGATCCTGAGGTAGA |
| D2 forward | CCCAGCAGAAGGAGAAGAAAG |
| D2R reverse | CAGGATGTGCGTGATGAAGA |
| GAPDH forward | GGAGAAACCTGCCAAGTATGA |
| GAPDH reverse | TCCTCAGTGTAGCCCCAAGA |

**Table S2**

| Antibody | #Catalogue | RRIDs | Dilution | Vendor |
|---|---|---|---|---|
| DAT | 22524-1-AP | AB_2879116 | 1:1000 | Proteintech |
| GAPDH | sc-32233 | AB_627679 | 1:2500 | Santa Cruz Biotechnology |
| GluA1 | ABN241 | AB_2721164 | 1:2000 | Millipore |
| GluA2 | AB1768-I | AB_2313802 | 1:1000 | Millipore |
| MAO-A | H00004128-B02P | AB_2137248 | 1:1500 | Novus |
| MAO-B | PA5-79624 | AB_2746739 | 1:1000 | Thermo Fisher Scientific |
| TH | MA5-38252 | AB_2898168 | 1:1000 | Thermo Fisher Scientific |
| Tubulin | 556321 | AB_396360 | 1:5000 | BD Biosciences |

**Table S3**

| Antibody | #Catalogue | RRIDs | Dilution | Vendor |
|---|---|---|---|---|
| Anti-mouse | ab216772 | AB_2857338 | 1:2500 | Abcam |
| Anti-rabbit | 611-144-002 | AB_1660962 | 1:2500 | Rockland |
